# Cannabidiol activates neuronal Kv7 channels

**DOI:** 10.1101/2021.08.20.457154

**Authors:** Zachary Niday, Laurel Heckman, Sooyeon Jo, Han-Xiong Bear Zhang, Akie Fujita, Jaehoon Shim, Roshan Pandey, Hoor Al Jandal, Selwyn Jayakar, Jennifer A. Smith, Clifford J. Woolf, Bruce P. Bean

**Affiliations:** Department of Neurobiology, Harvard Medical School, Boston, MA 02115; F.M. Kirby Neurobiology Research Center, Boston Children’s Hospital, Boston, MA 02115; ICCB-Longwood Screening Facility and Department of Immunology, Harvard Medical School, Boston MA 02115

**Author notes:** These authors contributed equally (Zachary Niday, Laurel Heckman, Sooyeon Jo).

**Keywords:** cannabinoids, Kv7.2, Kv7.3, M-current, sympathetic neuron, hippocampus, antiepileptic, antiseizure, Dravet Syndrome

## Abstract

Cannabidiol (CBD), a chemical found in the Cannabis sativa plant, is a clinically effective antiepileptic drug whose mechanism of action is unknown. Using a fluorescence-based thallium flux assay, we performed a large-scale screen and found enhancement of flux through heterologously-expressed human Kv7.2/7.3 channels by CBD. Using patch clamp recordings, we found that CBD at low concentrations activates Kv7.2/7.3 channels at subthreshold voltages, with 100 nM CBD producing a doubling of current at -50 mV. CBD shifted the voltage-dependence of channels in the hyperpolarizing direction, producing a shift in the midpoint of activation by ∼-14 mV at 300 nM. CBD also effectively enhanced native M-current in both mouse superior cervical ganglion neurons and rat hippocampal neurons. The potent enhancement of Kv2/7.3 channels by CBD seems likely to contribute to its effectiveness as an antiepileptic drug by reducing neuronal hyperexcitability.

## Introduction

Cannabidiol (CBD), a phytocannabinoid present in marijuana (Mechoulam et al., 1970), has been shown in recent clinical trials to be an effective agent for treating some forms of epilepsy, including Dravet Syndrome (Devinsky et al., 2017, 2018b, 2019; Miller et al., 2020) and Lennox-Gastaut syndrome (Devinsky et al., 2018a; Thiele et al., 2019). How CBD ameliorates epileptic activity is, though, unclear (Rosenberg et al., 2015,2017; Franco and Perucca, 2019). Unlike Δ(9)-tetrahydrocannabinol (THC), the other major phytocannabinoid in marijuana, CBD is not psychoactive and does not act as a direct primary ligand at CB1 or CB2 G-protein coupled receptors (Pertwee, 2005). At micromolar concentrations, CBD has inhibitory effects on a wide range of proteins, including many receptors and channels (Watkins, 2019). Like many classic antiepileptic agents, CBD inhibits voltage-dependent sodium channels in a state-dependent manner, with reported half-maximal concentrations of ∼2-4 10 μM (Hill et al., 2014; Patel et al, 2016; Ghovanloo et al. 2018; Mason and Cummins, 2020). However, as CBD reduction of overall epileptiform activity can be detected in brain slice preparations at concentrations as low as 100 nM (Jones, et al., 2010), the importance of sodium channel inhibition for CBD’s anti-convulsant effects remains uncertain (Hill et al., 2014). Other molecular targets that could mediate antiepileptic actions of CBD have been described, notably antagonism of the lipid-activated G protein-coupled receptor GPR55 (Ryberg et al., 2007; Kaplan et al., 2017), but so far electrophysiological effects correlated with GPR55 antagonism have been described only at concentrations of CBD ≥ 10 μM (Kaplan et al., 2017).

The most potent effect of CBD on electrophysiological function so far reported is an inhibition of endocannabinoid modulation of synaptic transmission (Straiker et al., 2018). This effect of CBD is mediated by a negative allosteric effect on CB1 receptors, with CBD acting at a site distinct from the primary binding site (Laprairie et al., 2015). Electrophysiologically, this inhibitory negative allosteric effect is detectable at 100 nM and is substantial at 500 nM (Straiker et al., 2018). Here we report that CBD acts at similar sub-micromolar concentrations to potently activate neuronal M-current, a non-inactivating potassium current mediated by Kv7 channels that activate at subthreshold voltages. CBD at concentrations as low as 100 nM shifts the voltage-dependence of activation of these channels in the hyperpolarizing direction, resulting in a significant activation of Kv7 current at subthreshold voltages. These results suggest that the activation of neuronal M-current is likely to be one mechanism by which CBD exerts its anti-epileptic action.

## Results

### CBD activates heterologously-expressed Kv7.2/7.3 channels

We discovered the ability of CBD to activate Kv7.2/7.3 channels in a screen using fluorescence signals from thallium entry evoked by depolarization of a CHO cell line stably expressing human Kv7.2 and Kv7.3 channels. In a screen of a library of 154 compounds chosen from structures with known or possible ion channel modulating activity, CBD was the only compound to produce a substantial enhancement of the fluorescence signal, except for retigabine and flupirtine, both known activators of Kv7.2/7.3 channels.

We then tested the action of CBD on the Kv7.2/7.3 cell line using whole-cell patch clamp recordings and saw a dramatic enhancement of the currents activated by depolarization, with particularly large effects for currents activated near -50 mV. Figure 1A shows an example, where 100 nM CBD produced a doubling of the current activated at -50 mV, while there was little effect at -20 mV, where channels are near-maximally activated in the control situation. 100 nM enhanced the current evoked at -50 mV by an average of 2.8 ± 0.4 (n=20), while 300 nM CBD enhanced the current by 4.6 ± 0.5 (n=17).

**Figure 1.**
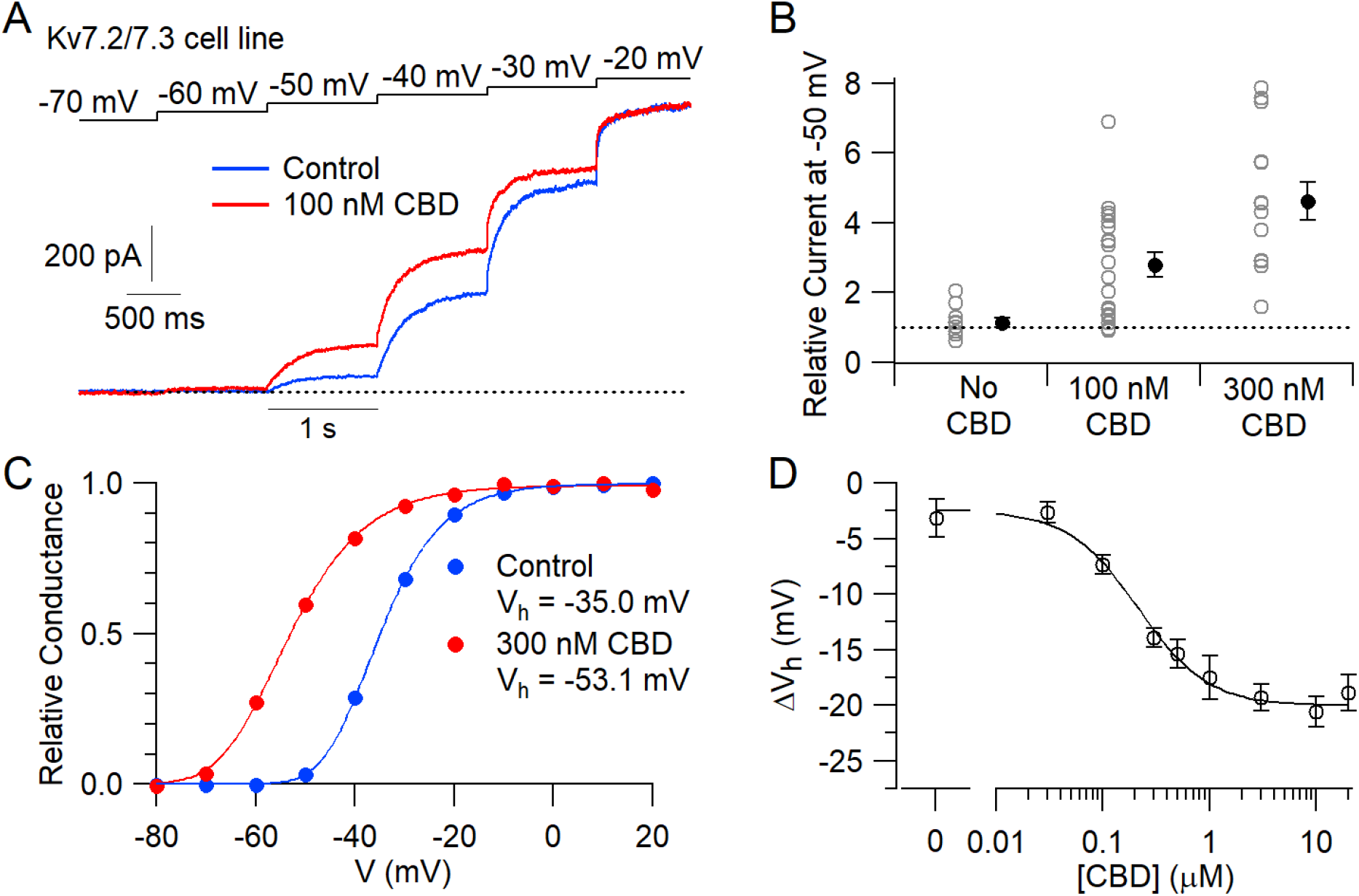
CBD enhancement of cloned human Kv7.2/7.3 channel current in CHO cells. (A) Whole-cell recording of hKv7.2/7.3 current evoked by staircase depolarizations before and after application of 100 nM CBD. (B) Collected results (mean ± SEM) for current evoked by a 1-s depolarization from -80 mV to -50 mV after a 6-minute exposure to 100 nM (n=20) or 300 nM CBD (n=14), normalized to current before CBD application. “No CBD” values (n=11) are for 6-minute dummy applications of solution containing only vehicle (DMSO). (C) Voltage-dependent activation of hKv7.2/7.3 channels measured in a cell before and after application of 300 nM CBD. Solid lines: Fits to data points of 4^th^ power Boltzmann function, [1/(1 + exp(-(V - V_hn_)/k))]^4^, where V is test pulse voltage, V_hn_ is voltage of half-maximal activation for single “n” particle, and k is slope factor for activation of n particles. Control: V_hn_ = -47.8 mV, k = 7.7 mV (midpoint of function= -35.0) ; 300 nM CBD: V_hn_ = -69.0 mV, k = 9.5 mV (midpoint of function -53.1 mV). (D) Concentration-dependent shift of activation midpoint by CBD. Measurements of the midpoint were made before and 10 minutes after exposure to CBD at various concentrations. mean ± SEM, n=9 for 30 nM CBD, n=21 for 100 nM CBD, n=17 for 300 nM CBD, n=12 for 500 nM CBD, n=7 for 1 uM CBD, n=15 for 3 uM CBD, n=19 for 10 uM CBD, n=10 for 20 uM CBD. Value for 0 CBD represents the measurement of a small shift that occurred with dummy applications of DMSO-containing control solution for 10 minutes (n=11). Solid line: fit to the Hill equation, ΔV_h_=-2.5 mV- 17.5 mV/(1+(EC_50_/[CBD])^n_H_), where EC_50_=213 nM and the Hill coefficient n_H_=1.3.

The enhancement of the Kv7.2/7.3-mediated current was produced by a shift of the voltage-dependent activation of the channels in the hyperpolarizing direction (Figure 1C). In collected results, 300 nM CBD shifted the midpoint for channel activation by an average of -13.9 ± 0.9 mV (n=17). The shift in the voltage-dependence of activation reached a maximum of about -20 mV at CBD concentrations of 3-10 μM, with CBD acting with a half-maximal concentration of about 200 nM (Figure 1D).

We next tested whether CBD enhances native Kv7 channels in neurons, using measurements of M-current in mouse superior cervical ganglion (SCG) neurons. Using the classic voltage protocol for distinguishing M-current from other potassium currents by virtue of its non-inactivating property and activation at subthreshold voltages (Brown and Adams, 1980), we used a steady holding voltage of -30 mV and hyperpolarizing voltage steps to quantify the M-current from its characteristic slow, voltage-dependent deactivation. Application of 300 nM CBD produced a dramatic enhancement of the steady-state outward current at -30 mV and also enhanced the slowly deactivating current seen during hyperpolarization to -70 mV (Figure 2A). It was also clear that CBD shifted the voltage-dependence of M-current activation, resulting in less complete deactivation for a step to -60 mV. In the collected results, 300 nM CBD produced an enhancement of steady current at -50 mV by a factor of 2.4 ± 0.3 (n=7).

**Fig. 2.**
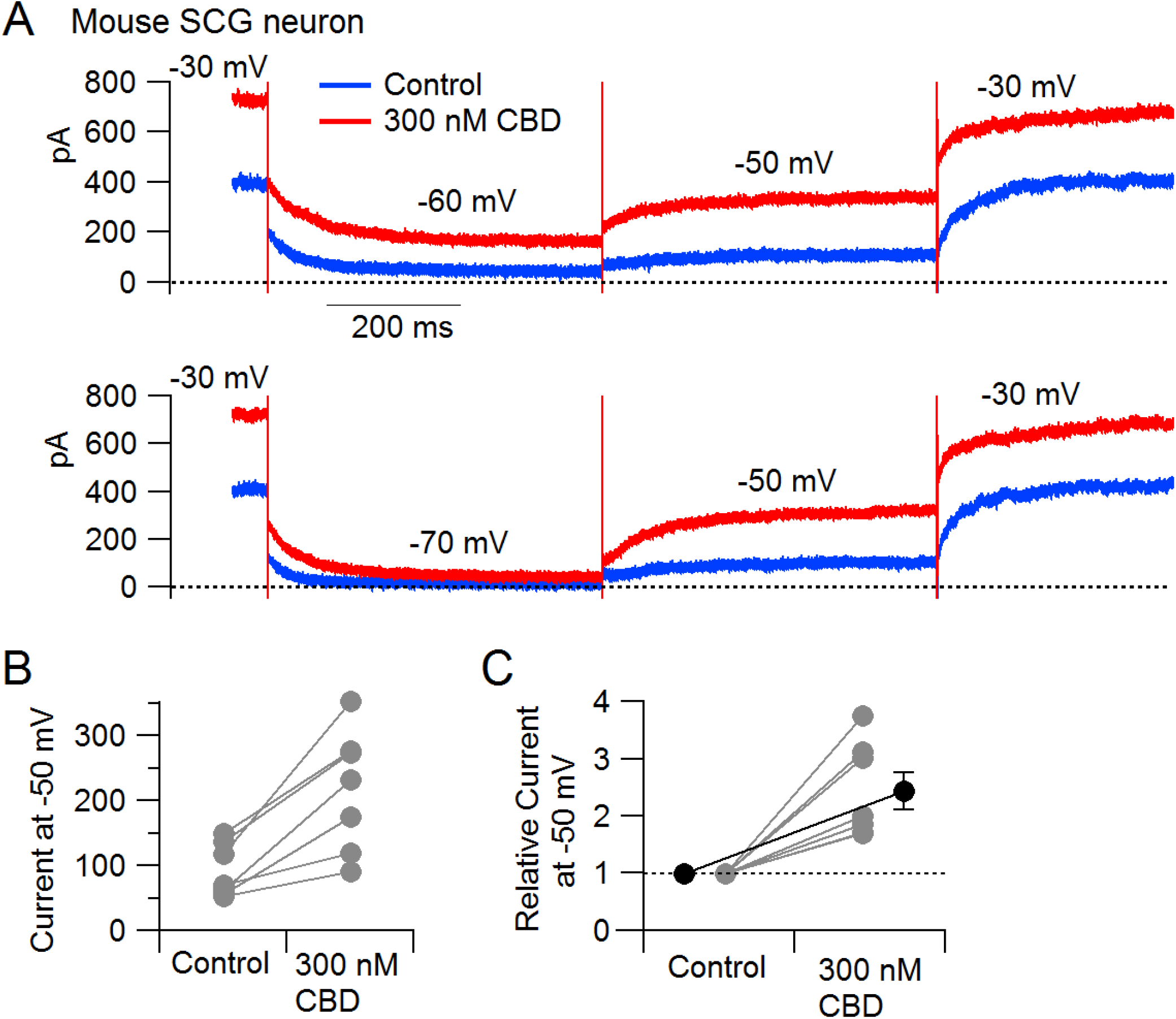
CBD enhancement of M current in mouse sympathetic neurons. (A) Currents evoked by hyperpolarizations to -60 mV and -70 mV from a holding potential of -30 mV before (blue) and after (red) application of 300 nM CBD. (B) Collected results for effect of 300 nM CBD on steady-state current at -50 mV (n=7). (C) Collected results for effects of 300 nM CBD on current at -50 mV, normalized to control current. Gray circles: individual cells. Black circles: mean ± SEM.

**Fig. 3.**
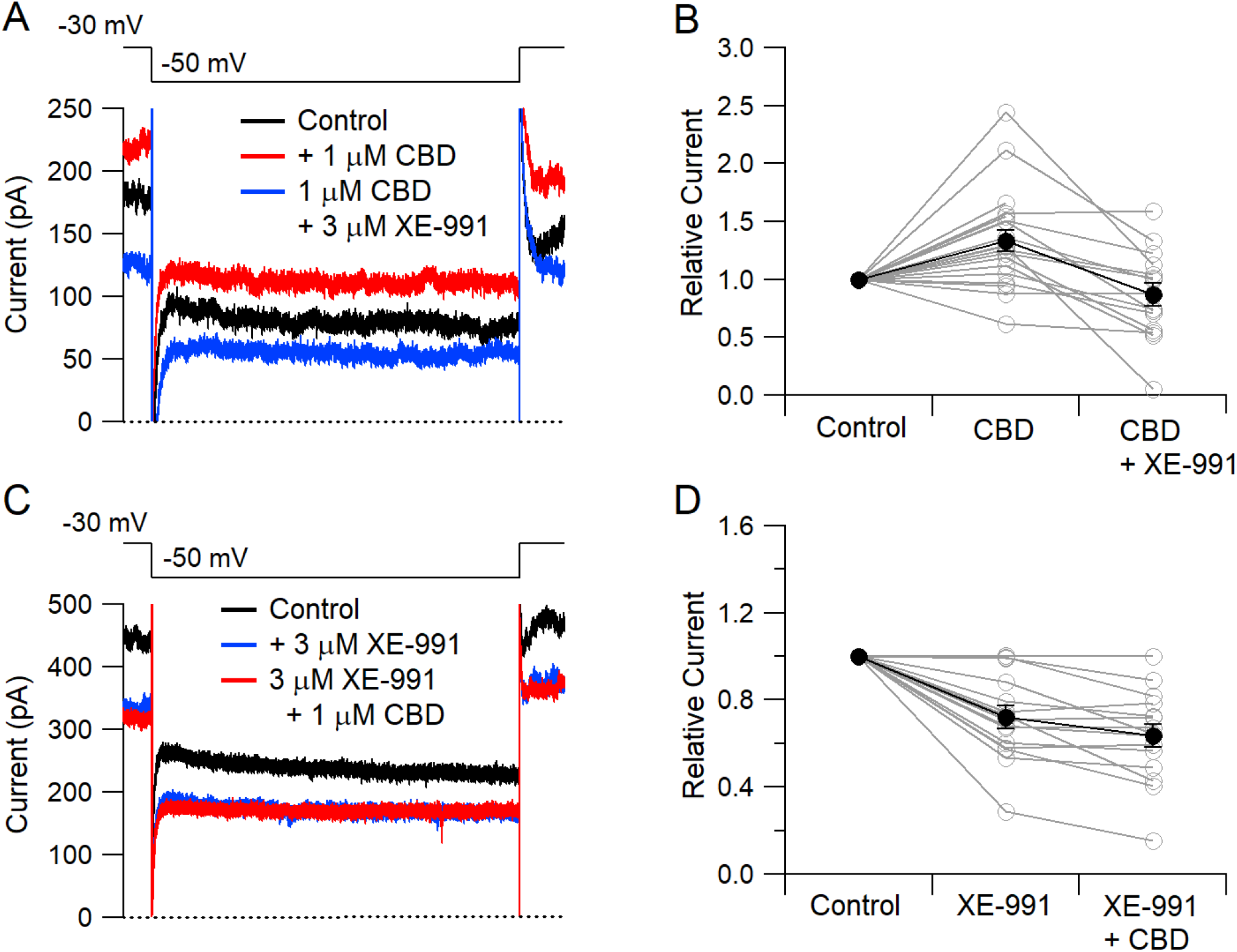
CBD enhancement of Kv7 current in rat hippocampal neurons. (A) Currents at a holding voltage of -30 mV and during a 500-ms hyperpolarization to -50 mV in control, after application of 1 μM CBD, and after addition of 3 μM XE-991 in the continuing presence of CBD. (B) Collected data with this protocol. Current was measured at the end of the step to -50 mV, normalized to current before application of CBD. Connected open circles indicate data for individual cells (n=20 for application of CBD, n=16 for application of CBD followed by XE-991) and closed circles represent mean ± SEM. (C) Currents in control, after application of 3 μM XE-991, and after addition of 1 μM CBD in the continuing presence of XE-991. (D) Collected data with symbols as in (B); n=15 cells for application of XE-991 followed by CBD.

To test whether CBD enhancement of M-current also occurs in central neurons likely involved in epilepsy, we tested CBD on potassium currents in hippocampal neurons. To facilitate application of well-defined concentrations of CBD without potential problems from absorption into the bulk tissue of brain slices, we used a preparation of cultured rat hippocampal neurons. Using a voltage protocol designed to emphasize M-current (holding the neurons at -30 mV and stepping to -50 mV), CBD enhanced the outward current at both -30 mV and -50 mV in 16 of the 20 cells tested. Consistent with this action of CBD being an enhancement of Kv7-mediated current, there was no increase if CBD was applied in the presence of the Kv7 inhibitor XE-991.

## Discussion

Kv7 channel-mediated M-current plays a major role in controlling the excitability of many types of neurons, including neocortical pyramidal neurons (Barrese et al., 2018; Brown and Passmore, 2009; Gunthorpe et al., 2012; Vigil et al., 2020). Enhancement of M-current is a clinically-proven mechanism of antiepileptic action, as demonstrated by the clinical efficacy of retigabine, an antiepileptic drug that acts by enhancement of current through Kv7 channels (Tatulian et al., 2001; Gunthorpe et al., 2012; Sills and Rogawski, 2020). Our results suggest that the clinical efficacy of CBD may also result largely or at least in part by the enhancement of the Kv7-mediated M-current in central neurons. As in the case of retigabine, it remains to be determined exactly which populations of neurons are most sensitive to this enhancement of M-current, and how these effects alter the overall network activity relevant to epileptic activity.

Interestingly, the effect of CBD in enhancing the neuronal M-current is the opposite to the effect of cannabinoids that act as agonists at the CB1 receptor, which inhibit M-current in hippocampal neurons (Schweitzer, 2000). Thus the fact that CBD is not a CB1 agonist - and actually acts as an allosteric antagonist at CB1 receptors (Laprairie et al., 2015; Straiker et al., 2018) - may be an important aspect of its mechanism of action. The opposite effects on M-current of CBD and CB1 agonists like THC, fit well with the history of the development of CBD as an anti-epileptic drug, which began with anecdotal evidence that extracts from a particular strain of cannabis with high CBD and low THC (“Charlotte’s Web”) were an effective adjunctive therapy for a child with Dravet Syndrome (Maa and Figi, 2014; Rosenberg et al., 2015; Williams and Stephens, 2020).

Our results add to recent experiments demonstrating that Kv7.2/7.3 channels are susceptible to enhancement by a wide variety of agents acting by several different mechanisms (Manville and Abbott, 2018a,b; Wang et al., 2018; Kanyo et al., 2020; Kurata, 2020). Such agents include endogenous compounds like GABA (Manville et al., 2018), the ketone body β-hydroxybutyrate (Manville et al., 2020), and arachidonic acid metabolites and derivatives (Schweitzer et al., 1990, 1993; Larsson et al., 2020a,b), as well as a variety of natural products including cilantro (Manville and Abbott, 2019). Further development of Kv7.2/7.3 enhancers for treating epilepsy and other neuronal disorders seems promising (Barrese et al., 2018; Vigil et al., 2020), especially because retigabine has been withdrawn from clinical use because of a number of off-target side effects (Brickel et al. 2020). Of all the classes of compounds found to enhance Kv7.2/7.3 channels, CBD has the unique distinction of having already been successfully used in multiple epilepsy clinical trials. However, CBD is far from a perfect drug (Sekar and Pack, 2019) as it requires large dosages and has a complex pharmacokinetic profile that limit its effective oral administration (Millar et al., 2019). Improved knowledge of CBD’s most important molecular targets, which our results suggest include neuronal Kv7 channels, should allow for the design of novel compounds that retain its key molecular actions but with improved pharmacokinetics and reduced off-target effects.

## Acknowledgements

Supported by the NIH (NS105076, NS36855, NS110860), DARPA (HR0011-19-2-0022), and the Charles R. Broderick III Phytocannabinoid Research Initiative.

## Author Contributions

Z.N., L.H., S.J., H-X.B.Z., A.F., J. Shim, R.P., H.A.J, J. Smith, B.P.B, and C.J.W. designed the experiments; Z.N., L.H., S.J., H-X.B.Z., A.F., J. Shim, R.P., H.A.J, and J. Smith conducted the experiments and analyzed the data; L.H., Z.N., S.J. H-X.B.Z., A.F., J.S. B.P.B, and C.J.W. wrote the paper.

## STAR Methods

### Thallium Flux Assay

#### Cell Culture

CHO cells co-expressing Kv7.2 and Kv7.3 (Mayflower Bioscience, BSYS-KV7.2/3-CHO-C) were cultured at 37ºC in 5% CO_2_ in a Thermo Scientific incubator in Ham’s F12-Glutamax-l medium (Gibco, Catalog # 31765-035) supplemented with 10% fetal bovine serum (Gibco), 1% penicillin/streptomycin solution (Gibco, Catalog # 15140-122) and 5 μg/mL puromycin (InVivogen, cat #ant-pr-1). Cells were seeded in 15 cm dishes at 200,000 cells per dish, fed twice weekly and cultivated once weekly. Twenty-four hours before the start of the screen, the culture dishes were trypsinized, and a Countess™ automated cell counter (Invitrogen) was used to quantify cell numbers before plating them into 4 Greiner poly-D-lysine-coated 384-well black clear-bottomed microplates at 20,000 cells per well in 40 μL media using a Multidroptm Combi Reagent Dispenser. The 4 microplates were incubated overnight in a Thermo Scientific incubator at 37ºC in 90% humidity and 5% CO_2_.

#### Compound Preparation and Handling

A custom library of 154 compounds oriented toward known or possible ion channel modulators was assembled, with each compound plated at 4 different concentrations in DMSO (0.08 mM, 0.4 mM, 2 mM, and 10 mM). Using a custom Seiko compound transfer workstation, 300 nL of experimental compounds, as well as positive (retigabine at 10 mM in DMSO) and negative (DMSO) controls, were pin transferred into a Greiner Bio-One 384 Deep Well Small Volumetm polypropylene microplate containing 30 uL of 1x FluxORtm chloride-free buffer. This resulted in 16 positive and 16 negative control wells on every assay plate and final compound concentrations between 267 nM and 33 μM. Each compound microplate was screened in duplicate.

#### Kv7.2/7.3 assay

The FluxORtm potassium channel assay (ThermoFisher) was performed using a Hamamatsu FDSS 7000 plate reader essentially as outlined in the product sheet. After the Kv7.2/7.3 CHO cells were incubated in the 384-well assay microplates for 24 hours, a 40 mL solution of FluxORtm dye was made by combining 400 μL Powerloadtm concentrate (100X), 40 μL of 13 FluxORtm II green reagent (1000X fluorescent dye) in DMSO, 31.2 mL purified water, 4 mL 10X FluxORtm assay buffer, 4 mL FluxORtm II background suppressor, and 400 μL probenecid (100X in water). Next, media was removed from each well of the assay microplates containing Kv7.2/7.3 CHO cells using an Agilent Bravo Liquid Handling system. The assay microplates were then washed two times with FluxORtm chloride-free buffer diluted from 5X to 1X (20 uL per well per wash). After the second wash was removed, 20 μL of the 40 mL dye solution was added to each 384-well assay microplate. The assay microplates were incubated in the dye solution at room temperature protected from light for 45 minutes. Subsequently, 10 μL of diluted compounds were added to each assay microplate from the compound dilution plate described above. Assay microplates were incubated in compound and dye for an additional 15 minutes at room temperature protected from light. Prior to beginning the assay, stimulus buffer was prepared by mixing 50 mM thallium sulfate (Tl_2_SO_4_, 4.8 mL), FluxORtm chloride-free buffer (5X, 6.0 mL), and purified water (19.2 mL). Next, 50 uL of this stimulus solution was loaded into an additional Greiner Bio-One 384 Deep Well Small Volumetm polypropylene microplate. The four assay plates and plate containing stimulus buffer were then loaded onto a Hamamatsu FDSS 7000Ex plate reader and liquid handler. For each cell microplate plate, 10 μL of stimulus buffer was added after 50 sec for a final concentration of 4 mM Tl+ in the assay plate. Fluorescence was measured for 600 data points (approx. 3 minutes) at 4 Hz. FDSSv3.3.1 software was used for baseline correction and data analysis.

#### Electrophysiology with CHO KV7.2/7.3 cell line

Cells were maintained and passaged in a humidified 37°C incubator in sterile culture flasks containing Ham’s F12-Glutamax-l medium (Gibco, Catalog # 31765-035) supplemented with 10% fetal bovine serum (Gibco), 1% penicillin/streptomycin solution (Gibco, Catalog # 15140-122) and 5 µg/mL puromycin (InVivogen, cat #ant-pr-1) and cells were passaged at a confluence of about 50-80%. For electrophysiological recordings, cells were seeded onto 12 mm cover slips (Fisherbrand, Catalog #12-545-80). Whole-cell patch clamp recordings were made using a Multiclamp 700B Amplifier (Molecular Devices). Electrodes were pulled from borosilicate capillaries (VWR International, Catalog # 53432-921) on a Sutter P-97 puller (Sutter Instruments) and shanks were wrapped with Parafilm (American National Can Company) to allow optimal series resistance compensation without oscillation. The resistances of the pipettes were 1.8-3.5 MΩ when filled with the intracellular solution consisting of 140 mm KCl, 10 mM NaCl, 2 mM MgCl_2_, 1 mm EGTA, 0.2 mm CaCl_2_, 10 mM HEPES, 14 mM creatine phosphate (Tris salt), 4 mM MgATP, and 0.3 mM GTP (Tris salt), pH adjusted to 7.4 with KOH. Seals were formed in Tyrode’s solution consisting of 155 mM NaCl, 3.5 mM KCl, 1.5 mM CaCl_2_, 1 mM MgCl_2_, 10 mM HEPES, 10 mM glucose, pH 7.4 adjusted with NaOH. After establishing whole-cell recording, cell capacitance was nulled and series resistance was partially (∼70%) compensated. The cell was then lifted and placed in front of an array of quartz fiber flow pipes (250 μm internal diameter, 350 μm external diameter, Polymicro Technologies, Catalog # TSG250350) attached with styrene butadiene glue (Amazing Goop, Eclectic Products) to a rectangular aluminum rod (cross section 1.5 cm × 0.5 cm) whose temperature was controlled by resistive heating elements and a feedback-controlled temperature controller (Warner Instruments, TC-344B). Solutions were changed (in ∼ 1 second) by moving the cell from one pipe to another. Recordings were made at 37 °C.

Voltage commands were delivered and current signals were recorded using a Digidata 1321A data acquisition system (Molecular Devices) controlled by pCLAMP 10.3 software (Molecular Devices). Current and voltage records were filtered at 5 kHz and digitized at 100 kHz. Analysis was performed with Igor Pro 6.12 (Wavemetrics, Lake Oswego, OR), using DataAccess (Bruxton Software) to import pClamp data.

The voltage-dependence of activation was measured from the initial tail current at a step to -50 mV following 1-s depolarizations to voltages between -100 mV and +40 mV from a holding potential of -80 mV. Current records were corrected for linear capacitative and leak current by subtracting scaled responses to signal-averaged 5 mV hyperpolarizations delivered from -80 mV. Tail current was averaged over a 1-msec interval starting at a time when the immediate jump in current had settled, typically 0.8-1.6 ms after the voltage step. Plots of normalized tail current versus test voltage could be fit well by a Boltzmann function raised to the 4^th^ power. The midpoint of activation was measured in a fit-independent manner by calculating the test voltage at which tail current reached half of its maximal value (reached at voltages between 0 to +40 mV), using linear interpolation between the test voltages straddling the midpoint. Calculation of shifts of activation midpoint by CBD was confined to cells in which maximal tail current at -50 mV remained at least 100 pA in the presence of CBD and in which the activation curve in CBD was fit well by a Boltzmann function raised to the 4^th^ power. Calculation of the enhancement of current at -50 mV was confined to cells in which the current was at least 20 pA in control.

CBD (Cayman Chemical, Catalog # 90080, CAS 13956-29-1) was prepared as a 10 mM stock solution in DMSO which was diluted in the external Tyrodes’s solution to the final concentration. DMSO was added to the control solution at the same concentration as in the CBD solution. In early experiments, CBD-containing solutions were prepared in polystyrene test tubes and applied to cells from reservoirs made from 10 mM polypropylene syringe bodies. Realizing that phytocannabinoids have exceptionally high lipophilicity (Thomas et al., 1990) and can apparently partition into plastic (Hippalgaonkar et al., 2011), we then switched to using glass reservoirs from which solutions flowed through hollow quartz fibers to be applied to cells. We found that using glass reservoirs and tubing resulted in larger and more reproducible effects of CBD concentrations of 1 μM and below. Reported data for these concentrations are confined to experiments using glass reservoirs and tubing. Effects of concentrations of 3 μM and above were not less when using plastic reservoirs, and collected data for concentrations of 3-20 μM include experiments done with both plastic and glass reservoirs.

#### Preparation of superior cervical ganglion (SCG) neurons

Superior cervical ganglia were removed from adult Swiss Webster mice of either sex (postnatal day 56), cut in half and treated for 20 minutes at 37°C with 20 U/ml papain (Worthington Biochemical, Catalog # LS003126) in a calcium- and magnesium-free (CMF) Hank’s buffer (Gibco, Catalog # 14170-112) containing 137 mM NaCl, 5.36 mM KCl, 0.33 mM Na_2_HP_4_, 0.44 mM KH_2_PO_4_, 4.2 mM NaHCO_3_, 5.55 mM glucose, 0.03 mM phenol red. Ganglia were then treated for 20 minutes at 37°C with 3 mg/ml collagenase (type I; Roche Diagnostics, Catalog # 10103586001) and 4 mg/ml dispase II (Roche Diagnostics, Catalog # 37045800) in CMF Hank’s buffer. Cells were dispersed by trituration with a fire-polished glass Pasteur pipette in a solution composed of two media combined in a 1:1 ratio: Leibovitz’s L-15 medium (Gibco, Catalog # 11415-064) supplemented with 5 mM HEPES, and DMEM/F12 medium (Gibco, Catalog # 11330-032) and plated onto coverslips. Then cells were incubated at 37°C (5% CO_2_) for 2 hours, after which Neurobasal medium (Gibco, Catalog # 10888-022) containing B-27 supplement (Gibco, Catalog # A3582801), and penicillin and streptomycin (Sigma-Aldrich, Catalog # P4333). Cells were stored at room temperature and used within 48 hours.

#### Electrophysiology with superior cervical ganglion (SCG) neurons

Whole-cell patch clamp recordings were made using a Multiclamp 700B Amplifier (Molecular Devices) interfaced to a Digidata 1321A data acquisition system (Molecular Devices) controlled by pCLAMP 10.3 software (Molecular Devices). Electrodes were 2-4 MΩ when filled with the intracellular solution consisting of 140 mM K aspartate, 13.5 mM NaCl, 1.6 mM MgCl_2_, 5 mM EGTA, 9 mM HEPES, 14 mM creatine phosphate (Tris salt), 4 mM MgATP, 0.3 mM Tris-GTP, pH 7.2 adjusted with KOH, with shanks wrapped with Parafilm to allow optimal series resistance compensation (70-80%). Seals were formed in Tyrode’s solution consisting of 155 mM NaCl, 3.5 mM KCl, 1.5 mM CaCl2, 1 mM MgCl2, 10 mM HEPES, 10 mM glucose, pH 7.4 adjusted with NaOH and cells were lifted in front of quartz fiber flow pipes attached to a temperature-controlled aluminum rod. M-current was recorded with external Tyrode’s solution containing 1 μM TTX and 10 μM CdCl_2_. Recordings were made at 37 °C.

Voltage commands were delivered and current signals were recorded using a Digidata 1321A data acquisition system (Molecular Devices) controlled by pCLAMP 10.3 software (Molecular Devices). Current and voltage records were filtered at 5 kHz and digitized at 100 kHz. Analysis was performed with Igor Pro 6.12 (Wavemetrics, Lake Oswego, OR), using DataAccess (Bruxton Software) to import pClamp data.

#### Preparation of rat hippocampal neurons

Primary cultures of hippocampal neurons were prepared from rat embryos (E19 to E20). Pregnant female Sprague-Dawley rats were anesthetized with isoflurane. The skin was washed with 70% ethanol, the peritoneal cavity was opened, and embryos were transferred into ice-cold preparation solution (Ca2+/Mg2+-free HBSS (Gibco, Catalog # 14170-112) with 5 mM HEPES (Gibco, Catalog # 15630-080) and 1 mM sodium pyruvate (Gibco, Catalog # 11360-070) in a 100 mm petri dish on ice. Heads and brains were sequentially dissected from embryos, with the ice-cold preparation solution exchanged during each step. Under a dissecting microscope, the meninges were stripped away from the cerebral hemispheres and dorsal hippocampi were dissected with a fine scissor. The hippocampal pieces were transferred into a pre-warmed preparation solution containing 37 U papain (Worthington, Catalog # LS003126), 5 mM L-cysteine (Sigma, Catalog # C7352), and 1080 U DNase I (Sigma, Catalog # DN-25), incubated at 37 °C for 15 minutes, and then washed 3 times with enzyme-free warmed preparation solution. The preparation solution was then exchanged for a titration medium (EMEM, ATCC, Catalog # 30-2003), 5 % FBS (Gibco, Catalog # 16140-071) and 1x penicillin/streptomycin (P/S, Gibco, Catalog # 15140-122), and the hippocampal pieces were titrated using Pasteur pipettes fire-polished to two different tip sizes. After determining cell density using a hematocytometer, a maintenance medium (Neurobasal media (Gibco, Catalog # 21103-049), 2% B27 (Gibco, Catalog # 17504-044), 5 mM Glutamine (Gibco cat # 25030-081), and 1x P/S) was added into cell suspension to make cell density of 1∼1.5 × 10^5^/ml. Five poly-D-lysine (Sigma, Catalog # P-7405)-coated coverslips (Fisherbrand, Catalog # 12-545-80) were placed in 35 mm dishes and 2∼3 × 10^5^ cells were plated in each 35 mm dish (≥ 4∼6 × 10^4^ cells/coverslip). Neurons were maintained for 13-17 days *in vitro* (DIV). Every 2∼3 days, half of the medium was removed from the 35 mm dishes and replaced with the same volume of the fresh maintenance solution.

All experiments using animals were performed according to an institutional IACUC-approved protocol.

#### Electrophysiology with rat hippocampal neurons

Recordings were made from neurons after 13 to 17 days *in vitro*. Neurons with three processes and a pyramidal shape were selected for recording. To avoid problems arising from absorption of CBD to plasticware, recordings were made in an all-glass chamber made by attaching a glass ring (18 mm outer diameter, 3 mm height, Thomas Scientific 6705R24) to a glass-bottom microwell dish (MatTek # P35G-1.5-20-C). Whole-cell recordings were obtained using patch pipettes with resistances of 2.2 to 2.5 MΩ when filled with the internal solution, consisting of 140 mM K-gluconate, 9 mM NaCl, 1.8 mM MgCl_2_, 0.09 mM EGTA, 9 mM HEPES, 14 mM creatine phosphate (Tris salt), 4 mM MgATP, and 0.3 mM Tris-GTP, pH adjusted to 7.2 with KOH. The shank of electrode was wrapped with Parafilm to allow optimal series resistance compensation. Seals were obtained and the whole-cell configuration established in Tyrode’s solution consisting of 155 NaCl, 3.5 KCl, 1.5 CaCl_2_, 1 MgCl_2_, 10 HEPES, 10 Glucose, pH adjusted to 7.4 with NaOH, with added 1 μM TTX. Reported membrane potentials are corrected for a liquid junction potential of -13 mV between the K-gluconate based internal solution and the Tyrode’s solution in which current was zeroed at the start of the experiment. The amplifier was tuned for partial compensation of series resistance (typically 40-70% of a total series resistance of 4-10 MΩ), and tuning was periodically re-adjusted during the experiment. Currents were recorded with a Multiclamp 700B Amplifier (Molecular Devices), filtered at 5 kHz with a low-pass Bessel filter, and digitized using a Digidata 1322A data acquisition interface controlled by pClamp9.2 software (Molecular Devices). Recordings were made at 30 °C.

M-current was evoked by 500-ms steps to -50 mV from a steady holding potential of -30 mV Stock solutions of 10 mM CBD in DMSO and 20 mM XE-991 in DMSO were made in glass vials and diluted into Tyrode’s solution (in glass vials) as 20 μM CBD or 60 μM XE-991 on the day of recording. Aliquots of these solutions were applied directly into the glass chamber and mixed with a 100 μL pipettor to make final concentrations of 1 μM CBD or 3 μM XE-991 respectively. To minimize any residual effect of CBD from the previous recording, the glass chamber was rinsed with 70% ethanol for 3 times and distilled water for 3 times before putting a new coverslip into the chamber.

